# Incentive value and spatial certainty combine additively to determine visual priorities

**DOI:** 10.1101/188045

**Authors:** K.G. Garner, H. Bowman, J.E. Raymond

**Affiliations:** School of Psychology, University of Birmingham

**Keywords:** attention, prediction, expectation, reward, incentive

## Abstract

How does the brain combine information predictive of the value of a visually guided task (incentive value) with information predictive of where task relevant stimuli may occur (spatial certainty)? Human behavioural evidence indicates that these two predictions may be combined additively to bias visual selection (additive hypothesis), whereas neuroeconomic studies posit that they may be multiplicatively combined (expected value hypothesis). We sought to adjudicate between these two alternatives. Participants viewed two coloured placeholders that specified the potential value of correctly identifying an imminent letter target if it appeared in that placeholder. Then, prior to the target’s presentation, an endogenous spatial cue was presented indicating the target’s more likely location. Spatial cues were parametrically manipulated with regards to the information gained (in bits). Across two experiments, performance was better for targets appearing in high versus low value placeholders and better when targets appeared in validly cued locations. Interestingly, as shown with a Bayesian model selection approach, these effects did not interact, clearly supporting the additive hypothesis. Even when conditions were adjusted to increase the optimality of a multiplicative operation, support for it remained. These findings refute recent theories that expected value computations are the singular mechanism driving the deployment of endogenous spatial attention. Instead, incentive value and spatial certainty seem to act independently to influence visual selection.

---

Humans are good at learning that specific sensory information, or cues, can predict subsequent events. For example, we learn quickly that hearing a siren on the left predicts a speeding emergency vehicle appearing from that direction, or that seeing a smile predicts a likely future opportunity to gain social approval. Knowledge about where and when new, important sensory information may appear or new reward opportunities may arise is only useful, however, if such knowledge can influence how cognitive mechanisms prioritise information representation for the eventual control of behaviour. Yet, our understanding of how learning and experience modify such prioritisation of visual signals, i.e. *visual selection*, remains incomplete. Particularly unclear is how multiple concurrent sensory cues, each associated with and therefore predictive of specific consequent outcomes, are combined to influence visual selection. A central tenet of many cognitive, computational, and neurobiological theories of visual selection (Buschman & Kastner, 2015; see Itti & Koch, 2001; Moore & Zirnsak, 2017, for reviews), is that competition for high-level neural representation of external objects is flexibly biased by information pertaining to goal-directed outcomes. Indeed, multiple endogenous sources appear to exert parallel influences on visual selection (see Hutchinson & Turk-Browne, 2012 for a review), even when learned information is antithetical to the current goals as defined by the task-set (see Awh et al., 2012 for a review). In the current study, we specifically seek to understand how selection biases that stem from learned associations between sensory cues and reward outcomes (incentive value) are combined with biases driven by associations between sensory cues and the probable location of task relevant target information (spatial certainty).

Recently, a large number of studies have shown that visual selection can be biased by the learned predictiveness and incentive value signalled by sensory cues (see Le Pelley et al., 2016 for a review). Such studies typically present stimuli in a simple selection task that probabilistically predict the reward magnitude available for correct performance, allowing those stimuli to become value associated. Then, in a subsequent visual selection task, these stimuli are presented as targets or distractors. The central finding is that visual selection performance is better or worse, respectively, compared to when such stimuli are absent (Anderson et al., 2011; Chelazzi et al., 2014; Raymond & O’Brien, 2009; although see Sha & Jiang, 2016). Convergent evidence from the behavioural (Le Pelley et al., 2016), neuro-cognitive (e.g. Barbaro et al., 2017; Hickey et al., 2010; Raymond & O’Brien, 2009) and neuro-economics (e.g. Dorris & Glimcher, 2004; Platt & Glimcher, 1999; Stanisor et al., 2013) literatures also provides a convincing picture that value cues influence visual selection proportionally to their associated reward value, underscoring the role of prior experience.

Another large literature arising much earlier (in the 1980’s) has shown that sensory signals that predict where task-relevant information might appear also bias visual selection (Posner 1980). When centrally presented, symbolic cues (e.g., arrows) are highly predictive of a target’s location (valid cues), response times (RTs) are faster and accuracy better compared to when spatial cues predict the wrong location (invalid cues). Central, symbolic spatial cues are often termed endogenous due to the assumption that their ability to bias visual selection largely stems from previously acquired and internally accessed knowledge. They are often contrasted with exogenous cues which are typically bright, brief, peripherally presented stimuli that appear at or very near probable target locations and are thought to modulate visual orienting in a largely reflexive manner. In the natural environment (and often in the lab), stimuli that predict where visual targets may appear probably activate a combination of endogenous biases and reflexive orienting mechanisms. However, our interest here is how endogenous (learned) spatial associations interact with other learned associations, specifically value associations to control visual selection. Strong evidence that spatial cueing effects (both endogenously and exogenously driven) depend on learning is that they scale with the reliability of the cue (Lanthier et al., 2015; Vossel et al., 2006, 2015), being stronger when cues are more predictive and weaker when predictability is low. Indeed, cueing effects appear to be a linear function of the certainty gained (in bits of information) by a spatial cue (Prinzmetal et al., 2015).

Although the issue of how multiple endogenous influences act on goal-directed visual selection has been previously addressed (Klink et al. 2014; Awh et al. 2012), it remains unclear how incentive value and spatial certainty information provided by pre-target cues are combined. One possibility, the *Expected Value Hypothesis*, is that visual selection is biased by the relative expected value, i.e., incentive value multiplied by spatial certainty for each outcome. Derived from economic theory (Morgenstern & Von Neumann, 1953), expected value is conventionally defined as reward magnitude multiplied by reward probability; in the context of selective attention in response to pre-target cues, such a computation would be the product of cue incentive value and spatial certainty. It is likely to be normalised across potential outcomes given the trial context (Padoa-Schioppa & Assad, 2008). Evidence for such an operation would suggest that incentive value and spatial certainty are combined by a common, perhaps singular mechanism to exert influence on visual selection. Support for this theory comes from the finding that human saccadic initiation times are more tightly correlated with the relative expected value of potential upcoming target locations than with either the spatial certainty for the target location, or its incentive value alone (Milstein & Dorris, 2007). Moreover, close correspondence has been shown between neurons that fire proportionally to the incentive values associated to sensory cues and those that change their response to correlate with the spatial certainty introduced by a directional cue, at least in macaque V1 (Stanisor et al., 2013). Indeed, this latter finding was interpreted as providing evidence for a singular neural mechanism able to re-weight incentive values across the visual scene on-the-fly by updating computation of expected values, given post-cue spatial probabilities.

A second alternative is the Additive Hypothesis that posits incentive value and spatial certainty to exert independent, as opposed to multiplicative, influences on visual selection. Origins for this idea stem from computational theories proposing that variation in physical salience within a scene (a saliency map) for different stimulus dimensions (e.g., colour, motion or orientation) are independently calculated, then weighted and summed to create a visual priority map (Zhao & Koch, 2012). Indeed, empirical investigations support the notion of additive effects for the combination of salience information based on different features, and report that nonlinear combinations across feature dimensions arise only when there is overlap between the underlying saliency mechanisms (Nothdurft, 2000). Although suggestive of the possibility that incentive value and spatial certainty might combine additively to control selection, it remains unclear whether additivity could apply to non-physical features, such as learned associations.

However, preliminary support for this possibility has been reported by Stankevich and Geng (2014). They asked participants to detect as quickly and accurately as possible a simple target that could appear on the left or right within a coloured placeholder. In their experiment, placeholder colour indicated the magnitude of response-contingent rewards, as it does in the experiment we report here. However, they provided no explicit spatial cues as to target location. Instead, across blocks and without instruction, the probability of the target appearing on one side versus the other was varied. Greater performance benefits were observed when the target appeared in the more probable location and when that location corresponded to a high versus low value placeholder. Critically, these benefits were additive, suggesting that incentive value and spatial certainty acted independently to influence visual selection, according to additive-factors logic (Sternberg, 1998). However, in their experiment, spatial-certainty was established over many trials, allowing predictions about target location to be generated well before each trial began and certainly in advance of location-specific incentive information. This may explain why incentive information provided an additive “top-up” effect. Such effects might not occur when location-specific incentive information is available first and spatial certainty cues providing task relevant information are presented subsequently. Accordingly, how two different endogenous cues (incentive and spatial) might be combined in humans when they are available only via new sensory information as each trial unfolds remains unknown.

To address this issue, we conducted two experiments that combined the methods of relevant previous studies to directly test these two hypotheses. As described in Figure 2, on each trial, participants viewed two spatially separated coloured placeholders that served as incentive cues for 400-500 ms; placeholder colour signalled the reward value for a subsequently presented letter target, should it appear in that placeholder and be correctly identified. Then, a central symbolic spatial cue was presented that signalled which placeholder would more likely encircle the upcoming target. Then, 400 ms after spatial cue onset, one letter briefly appeared in each placeholder. The task was to identify the target (as one of two possible letters) as fast and accurately as possible. Incentive value for the potential target location was always either high or low on every trial; however, the reliability of the spatial cue (spatial certainty) was varied between blocks. The Expected Value Hypothesis, implicating a multiplicative relationship between incentive value and spatial certainty, predicts an interaction of these two variables on performance. See Figure 1. Specifically, it predicts a super-additive effect when spatial certainty is high and a sub-additive influence when it is low. In contrast, the Additive Hypothesis implies simple additivity of incentive value and spatial certainty. Evidence for this would be simple main effects on performance of incentive value and spatial certainty with no interaction effect. In Experiment 1, we varied spatial certainty over a wide range. In Experiment 2, rewards for correct responses diminished as RT lengthened, creating conditions favouring the use of an expected value mode of cue combination. Results were evaluated using a Bayesian model selection approach. To anticipate, we found in both Experiments that the influence of incentive value and spatial certainty remains additive across all tested levels of spatial certainty, arguing against the expected value hypothesis.

**Figure 1:**
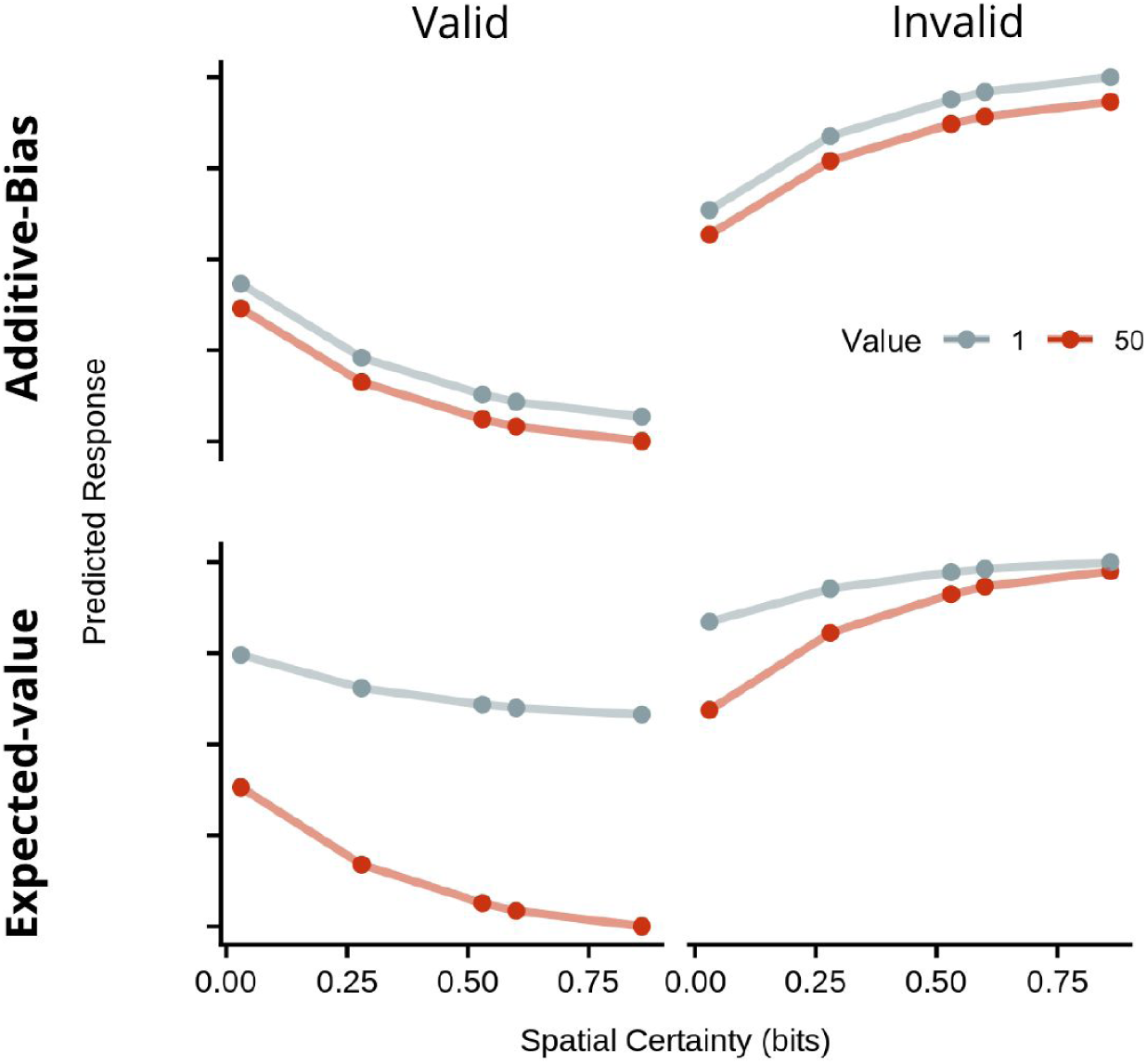
Illustration of theoretical predictions. Predicted RTs in arbitrary units, plotted by cue validity (columns), spatial certainty (sc, x-axis) and incentive value (iv, colours), according to an additive bias operation (y ∼ -β1sc + -β2iv) or an expected-value model (y ∼ -βsc*iv). (Note: the spatial certainties used in this simulation are the same as those chosen for Experiment 1).

**Figure 2:**
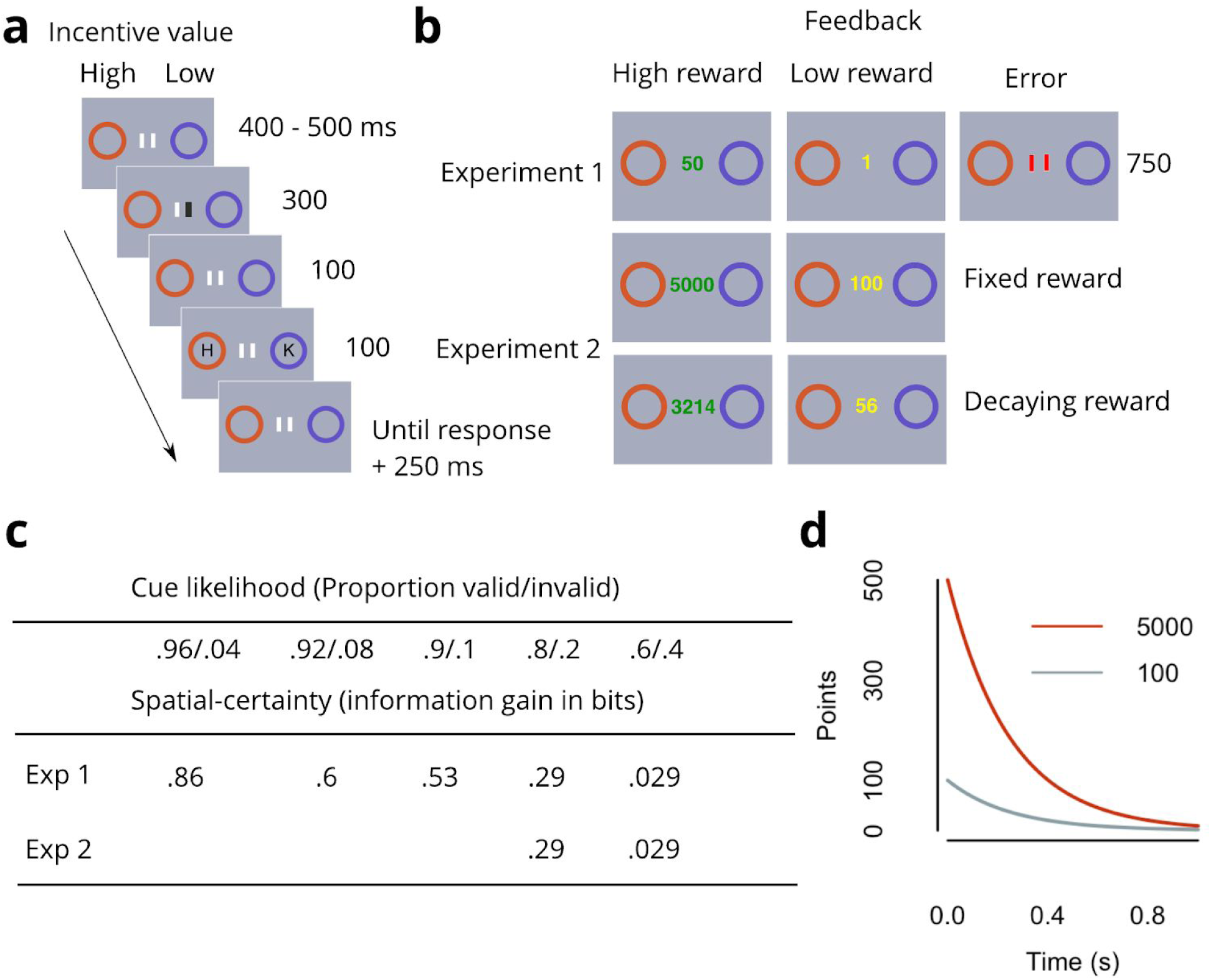
Study method. Task procedure and feedback conditions for Experiment 1 and Experiment 2. **a**. Trial structure: Participants monitored two different coloured circular placeholders (incentive cues). Colour indicated the magnitude of a performance-contingent reward for correct target (“H” or “N”) identification, should the target subsequently appear within that placeholder. Then, one of two central bars darkened, indicating the more likely target location (left, right). **b**. Reward feedback structure: After response + 250 ms, performance feedback and response-contingent rewards were presented as shown. In Experiment 2, reward feedback was either independent of response time (fixed) or decremented exponentially after target onset until response (decaying). **c**. Spatial certainty was parametrically manipulated across blocks by increasing the information gained (in bits) from the spatial cue. **d**. Logic of the decaying reward condition in Experiment 2. Figure shows reward value available as a function of time from target onset. As the expected value computation involves a multiplicative weighting of spatial certainty and incentive value, responses should be super-additive or sub-additive depending on the spatial certainty/incentive value combination. As response times should reflect the inverse of this weighting, responses should be faster in a high certainty/high incentive scenario than responses based on an additive operation, and slower in a low certainty/low incentive scenario than an additive operation. Applying an exponential decay function to the incentive value at target onset means that the extra rewards accrued by being faster towards high incentive value cues (change in the high (5000) value on the y-axis, while moving leftward on the x-axis) would outweigh the losses accrued from being slower towards low incentive value cues (change in the low (100) value on the y-axis, while moving rightward on the x-axis). Therefore, any operation that favours this response pattern would accrue greater total rewards than an additive operation, and therefore may emerge under such reward conditions.

## Experiment 1

Five levels of spatial certainty were tested. Although it is conventional in attention cueing experiment to use valid spatial cues on 80% and invalid cues on 20% of trials, we included conditions wherein the computation of spatial certainty is trivial, i.e., as the spatial cue approaches 100% validity. We reasoned that if, as the Expected Value Hypothesis posits, a single mechanism responds to both incentive value and spatial certainty information, then more resources would be available to respond to value information under high versus low spatial certainty conditions, leading to greater versus lesser effects of incentive value, thereby showing super-additivity. To test this idea, we more densely sampled valid trial probabilities between .90 and .96 as well as the more conventional probabilities of .8 and .6.

## Method

All the task and analysis code, and data from the current study are available online^1^. The trial sequence of the spatial-orienting task (Posner, 1980) used to assess the combined influence of spatial certainty and location specific incentive cues, and the key manipulations for Experiments 1 and 2, are shown in Figure 2.

## Participants

As larger samples protect against spurious findings (Button et al., 2013; Lorca-Puls et al., 2018), we opted to double the sample size of previous work correlating human performance with expected-value (N=10) (Milstein & Dorris, 2007), and recruit a minimum of 20 participants. We calculated the stopping rule for data collection as the number of weeks where testing at maximum capacity would bring us to at least the minimum sample size (6 weeks with 4 people per week, allowing recruitment of >20 participants in order to protect against dropouts). Participants were recruited if they were aged 18 years or over and reported normal or corrected-to-normal vision, with no history of psychiatric or neurological illness, injury, or disorder. Participants earned either course credit or payment (£7 per session), and any additional rewards accrued during the session (∼£10). All procedures were approved by the University of Birmingham Human Research Ethics Committee. A total of 23 participants were recruited. Of these, 1 was excluded due to technical failure and a second due to experimenter error. The remaining 21 participants (19 female, 18 right-handed, mean age = 20.3, sd 4.5) completed all the procedures.

## Apparatus

All experimental procedures took place in a room with a single testing station, under conditions of low ambient light. All tasks were programmed in Matlab (Mathworks, Natick, MA, 2013a), using the Psychophysics Toolbox extension (Brainard, 1997; Pelli, 1997). The tasks were run on a Stone SOFREP-144 computer with a 23-inch Asus VG278HE monitor (1920 × 1080 pixels, 60-Hz refresh) viewed from 57 cm.

## Stimuli

Two white [RGB: 200, 200, 200] vertical lines (0.5° w x 1° h) were presented in the centre of the screen. A darkening of one line [50, 50, 50] served as the endogenous spatial cue. Comparable cues have been used in previous work examining the influence of dynamic cue reliabilities on the deployment of spatial attention (Vossel et al., 2015), and therefore seemed appropriate for the aims of the current study. Although this type of central spatial cue had a small lateralised element, possibly activating exogenous orienting mechanisms to some extent, they were centrally presented (with 3.75° between the outer edge of the central cue and the peripheral target), and critically, require some interpretation from the brain to map the relationship between the cue and target locations (Berger et al., 2005; Briand & Klein, 1987; Posner, 1980). Thus, the main characteristics of the cues used here resemble those used by previous researchers to study spatial certainty effects. Two coloured discs (2.2° deg in diameter), one in purple [87, 75, 80], the other in orange [120, 86, 1] (matched for luminance) served as value cues. They were aligned along the horizontal meridian and positioned 4.5° from the centre. Targets (‘H’ or ‘N’) and distractors [‘Z’ or ‘K’] were presented in light grey [90, 90, 90] Helvetica font, encompassed 1°, and were centred on the disc’s centre. Feedback was presented in green [0, 255, 0] for high reward values, amber [255, 191, 0] for low reward values, and red [255, 0, 0] for errors. All stimuli were presented on a grey [RGB: 118, 118, 118] background.

## Procedure

As shown in Figure 2, each trial began with the simultaneous presentation of both incentive value cues and two centrally presented vertical lines. After a pseudo-randomly chosen duration of 400-500 ms, the left or right fixation line darkened for 300 ms. After a further 100 ms, the target and distractor were presented for 100 ms (target identity was equiprobable for each incentive value x spatial certainty x cue validity condition). Participants pressed with the ‘v’ or ‘g’ key on a standard keyboard to indicate the target identity. After 500 ms, feedback was presented for 750 ms; either the central fixation was replaced with the high reward value, the low reward value, an error signal (fixation lines turned red), or the fixation remained the same (no reward). Rewards were awarded on 80% of correct trials to prevent feedback signals becoming redundant. The high and low incentive value cue locations and the target location were equally likely to be on the left or right; all conditions (cue value location, target location, and target identity) were fully crossed within each session. Colour/value pairings (e.g. purple = 50 points/orange = 1 point) as well as target-response mappings were counterbalanced across participants.

Across blocks, the likelihood of cue validity was varied to be either .6 valid/.4 invalid, .8/.2, .9/.1, .92/.08, .96/.04, resulting in information gains (spatial-certainty) of .029, .29, .53, .6 and .86 bits. Each block contained 100 trials. At each of 4 sessions, participants completed 4 blocks for each level of spatial certainty. Participants took between 4 days and 1.5 weeks to complete the experiment (block order was pseudo-randomised for each session). Participants completed 4 sessions so that we could have sufficient trial numbers to obtain a reliable estimate of performance for the low spatial certainty conditions. Target-value contingencies were split equally within each set of valid and invalid trials for each cue-likelihood condition.

Participants were explicitly instructed how many points were available should the target appear in the location of the high and low incentive value cues (50 vs 1 point), and were instructed that the incentive cues signified that points were available most, but not all of the time. They were also instructed that the darkened line was a clue to where the target could appear, however, they were not explicitly informed that the spatial cue’s reliability might vary. Participants were requested to keep their eyes at fixation, and to respond as accurately and as quickly as possible to the target. Participants were also informed that their points would be exchanged for cash at the end of the session (1000 = £1). At the start of the first session, participants practiced until they achieved at least 16/20 correct responses.

## Statistical Approach

### Data pre-processing

All data was analysed using the R programming language (v3.3.2) (R Core Team, 2013), and R Studio (v1.0.44) (RStudio Team, 2016). RT data were rejected if they were greater than +/- 2.5 s.d. from the mean for that participant in that condition. As participants were not explicitly informed when there was a change in spatial-certainty, we assumed that trials immediately subsequent to changes in spatial-certainty would be contaminated by learning effects. To remove the contaminated trials for each participant, we collapsed the data across spatial certainty blocks, and ordered the data according to trial number. We then fit piecewise linear regressions to find the break point that minimized the mean square error (MSE). Trials occurring prior to the breakpoint were removed (mean = 12.3, sd 8.0). However, when we performed the analyses without removing these trials, the pattern of results was the same.

### Model specification and selection

The aim of the study was to compare whether an additive model remained the best, given the data, even under conditions where an additive relationship could be expected to break down. The key aim of each analysis was to determine whether a model that included an interaction term between incentive value and spatial certainty was more probable, given the data, than one that only included main effects (i.e., an additive model). To quantify evidence, we used a Bayesian approach that provides the advantage of offering the ability to quantify evidence against a specific model, which is not possible using null hypothesis significance testing approaches (Nickerson, 2000; Wagenmakers, 2007). Additionally, Bayesian approaches protect against the problem of model complexity: although more complex models may predict with high likelihood a greater range of values, if these predictions are uninformative, this will result in a more diffuse marginal likelihood distribution when integrating across prior distributions for the parameters, thereby penalising the resulting model evidence. First, we fit all possible linear mixed models on the RT data, with the regressors being (a) incentive value of the target location, (b) cue validity, and (c) spatial certainty offered by the cue. Spatial certainty was computed in line with Prinzmetal et al (Prinzmetal et al., 2015). Specifically, Shannon’s (Shannon, 1948) measure of entropy (H) measures the amount of uncertainty in a probability distribution and is at maximum when the cue is completely unpredictable with regard to the target location. Therefore, spatial certainty gained by an informative cue can be calculated as:

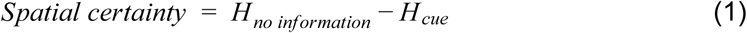

when *H* is defined in the standard manner:

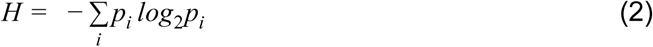

and *p*_*i*_ is the probability that the target appears at location *i*, given the cue. For example, with 2 locations, and a cue that is .8/.2 valid/invalid:

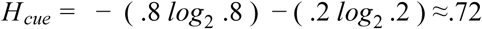

As *H*_*no information*_ is 1 (corresponding to complete uncertainty, i.e. .5/.5), then the information gained by the cue is 1 - .72 ≈ .28 bits.

After fitting all possible models to the RT data obtained in each experiment, we computed Bayes Factors (BFs) to quantify evidence for each linear mixed effects model against the null model (intercept plus random effects of participant) using the Bayes Factor package (Morey et al., 2014), and implementing the default Jeffreys-Zeller-Siow (JZS) prior on slope estimates (Liang et al., 2008). We then identified the six best performing models. We report the BF of the winning model relative to the null model, and the BF ratios between the best model and the next five best models, to demonstrate the strength of evidence in favour of the winning model. We follow the guidelines of Kass and Rafferty (Kass & Raftery, 1995) when interpreting the strength of evidence. This was sufficient to determine whether the evidence favoured a model that included only main effects, or an incentive value x spatial certainty interaction. All BFs are reported along with the proportional error of the estimate. For readers interested in confirming that a null hypothesis significance testing (NHST) approach yields the same conclusions, please refer to the online repository for this study.

### Accuracy data

Accuracy data were analysed to ensure the results were not due to a speed-accuracy trade-off. For each experiment, we fit all possible linear mixed models to the accuracy data, and selected the winning model. We then report the fixed effects estimates for each relevant factor in the winning model.

## Results

### RT

Group mean RT data (dots) and winning model fit (lines) are presented in Figure 3a. The main effect of incentive value was to speed RT by approximately 50 ms ± 3 (SE) for high versus low incentives. Spatial certainty served to increase the effect of cue validity; the difference between valid and invalid trials increased by approximately 90 ms ±29 (SE) across levels of certainty. Arguing against the expected value hypothesis, the preferred model included only main effects of each factor (incentive value, spatial certainty, and cue validity), and a spatial certainty x cue validity interaction term (BF, relative to the intercept only null model, = 1.74E+58, ± .87 %, see Figure 3b). Importantly, there was positive evidence that this model was preferred over the next best model (BF = 3.8, ± 1.45%), which was identical to the winning model except that it also included an incentive value x spatial certainty interaction term. Therefore, the evidence favours a model that does not include an interaction between incentive value and spatial certainty.

**Figure 3:**
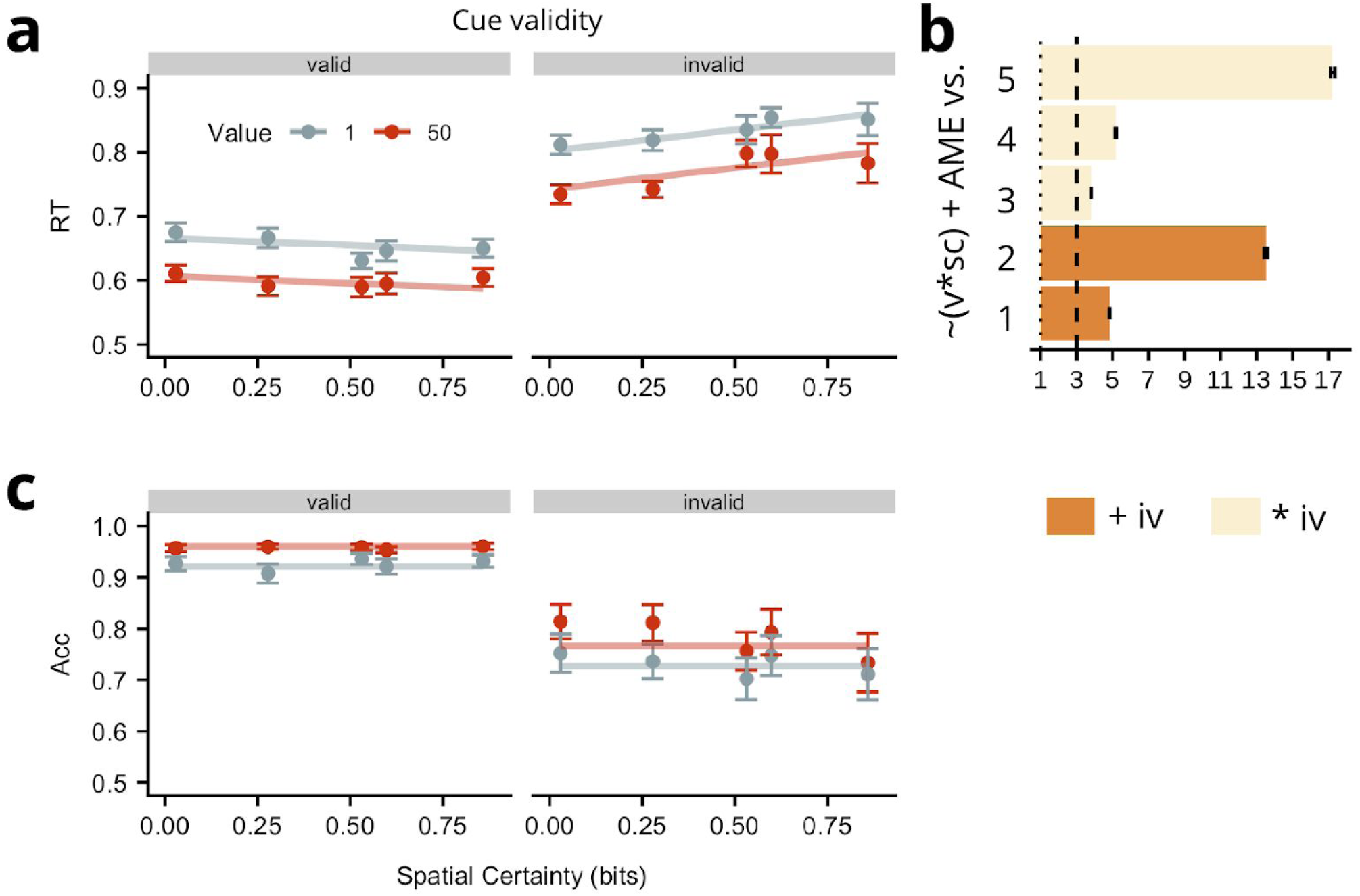
Results from Experiment 1. **a**. Observed group mean RTs in ms for targets appearing in high (dark circles) or low (light circles) value placeholders plotted as a function of spatial certainty (x-axis) for valid and invalid spatial cues (panels). Vertical lines indicate ± 1 within-subject s.e. Lines represent fit of the winning model. The winning RT model did not involve any interaction of incentive value (iv) and spatial certainty (sc) supporting an additive hypothesis, although it did indicate an interaction of sc and validity (v). **b**. BFs for the probability of the winning RT model (P[Win]: (v * sc) + AME) relative to the 5 next best models (Alternative, P[Alt], models, y-axis*). The larger the BF value, the stronger the evidence for the winning model*. Any values lower than 1 (dotted line) support P[Alt]; BF values between 3 (dashed line) and 10 constitute moderate evidence for the winning model, values > 10 provide strong evidence. Dark bars indicate P[Alt] contains only an additive influence of incentive value; light bars indicate P[Alt] involves a multiplicative influence of incentive value and either spatial certainty or validity. The Alt RT models were as follows: 1) ∼v + iv, 2) ∼AME, 3) ∼(v * sc) + (sc * iv) + AME, 4) ∼(v * sc) + (v * iv) + AME, 5) ∼(v * sc) + (v * iv) + (sc * iv) + AME. **c**. Observed group mean accuracy plotted as in panel a. Importantly, the accuracy data show that the RT findings were not due to a speed-accuracy trade-off. BF = Bayes Factor. v = cue-validity, sc = spatial certainty, iv = incentive value, rc = reward condition, AME = all main effects.

### Accuracy

The probability of an accurate response on invalid trials was .16 ±.02 (SE) lower than on valid trials and was .05 ±.02 (SE) less for targets appearing in low versus high incentive value cues. This shows that as participants became slower on invalid and low incentive value trials, they also became less accurate, arguing against the notion that effects were due to a speed accuracy trade-off (see Figure 3c). The preferred model for these data, relative to the null model included only main effects of incentive value and cue validity (BF = 8.83E+47, ±.56 %, relative to the null model).

## Discussion

There are two noteworthy findings from Experiment 1. First, even though the optimal strategy in this experiment would be to ignore the spatial cues entirely, especially in blocks with low spatial certainty, clear effects of valid versus invalid cues were found. Second, models posing an additive influence of incentive value and spatial certainty outperform those allowing an interaction between the two. This goes against the Expected Value Hypothesis and suggests that visual selection mechanisms do not integrate incentive value and spatial certainty, even when approaching the limits of certainty. Nevertheless, the second-best model to account for the RT data included an interaction between spatial certainty and incentive value, suggesting that this interaction is not entirely implausible. Perhaps additive effects would fail to provide the best description of the data if another form of appropriate pressure was applied to visual selection. Indeed, previous studies show that learning can direct the sampling of sensory information to optimise reward accrual (Drugowitsch et al., 2015; Kiani & Shadlen, 2009; Serences, 2008), allowing the possibility that a reward structure favouring an expected value operation may be sufficient to modulate the additive influence of incentive value and spatial certainty. The aim of Experiment 2 was to provide this test.

## Experiment 2

The aim of Experiment 2 was to test whether the additive influence found in Experiment 1 would persist even when the reward structure of the task renders additivity to be suboptimal. In Experiment 2, we leveraged the predictions made by the Additive and Expected Value operations to construct such a reward structure. As mentioned above, expected value computations are multiplicative, and should produce super-additive effects when both incentive values and spatial certainty are high, and sub-additive effects when incentive value and spatial certainty are low. Thus, RTs driven by an expected value operation should be faster than those driven by an additive operation when spatial certainty and incentive value are high and slower than an additive operation when spatial certainty and incentive value are low. Therefore, conditions that preferentially reward fast RTs on high-incentive, high-certainty trials and outweigh the costs incurred for slow RTs on low-incentive-low-certainty trials should favour adoption of an expected value strategy over an additive strategy (see Figure 2d). To produce these reward conditions, in Experiment 2, participants completed the same task as in Experiment 1 (albeit sampling fewer levels of spatial certainty), with an added condition wherein reward value exponentially decayed after target onset^2^. Of course, the assumption that these reward conditions favour a multiplicative (i.e. Expected Value) operation only holds if both spatial and incentive cues are leveraged to bias information processing. For example, an alternate and arguably optimal strategy would be to ignore the spatial cues entirely and to attend only to the high value location. However, given that we know from Experiment 1 that both cues effectively influence performance, making safe the assumption that decaying reward conditions favour a multiplicative over an additive operation.

Replicating the finding from Experiment 1, evidence supports the additive hypothesis but not the expected value hypothesis.

## Method

### Participants

We calculated the stopping rule for data collection as the number of weeks where testing at maximum capacity would bring us over the minimal sample size (3 weeks with 10 people per week). Of the 28 participants recruited, 1 was excluded due to technical difficulties with the eyetracker. A second participant was excluded as they did not meet the criterion required to terminate the practice. The remaining 26 (mean age = 19.5 years, sd = 1.03, 24 females, 26 right-handed) completed all the study procedures. Two of these participants had also completed Experiment 1.

### Apparatus

In addition to that used for Experiment 1, an Eyelink® 1000 desktop-mounted eye-tracker (SR Research Ltd., Ottawa, Ontario, Canada) recorded movements of the left eye with a sampling frequency of 500 Hz. This was used to ensure that eye movement were not contributing to results, even though participants were clearly instructed not to move their eyes (replicating instructions used in Experiment 1 that were not, however, verified for compliance).

### Stimuli

The stimuli were the same as in Experiment 1, except that the value cues were presented at 5.7°. This change was made to match the exact layout used in previous work (Stankevich & Geng, 2014).

### Procedure

The procedure was the same as Experiment 1 with the following exceptions. Participants’ eyes were monitored on every trial. If the participant’s eyes moved more than 50 pixels from the fixation at cue-offset, text appeared to notify participants they had been “too-fast”. The trial was then terminated. Terminated trials accounted for ∼3 % of all trials.

Cue-values were increased from Experiment 1 to 5000 vs 100 points, so that participants could gain at least 1 point when a decay was applied to the low incentive value. Correct responses were rewarded on 80% of trials as in Experiment 1. Given that there were 200 trials per condition (see below), and assuming ∼85% accuracy, this provides ∼136 rewarded trials per condition. Previous studies suggest that participants can detect dynamics in incentive value associations with far fewer trials (N=8, Ittipuripat et al, 2015) or lower reward-to-non-rewarded ratios (33:77%, Serences, 2008), and that these detections influence visual spatial priorities. In the decay reward-condition, an exponential decay function (reward value = points*(*e*-^*4*RT*^), RT = Response Time) was applied to each value at target onset. The monetary value of points was adjusted so that participants received the same rate of cash payments as Experiment 1 (100,000 = £1). Participants were informed at the start of the decay blocks that the value available to them would begin to run out upon appearance of the target.

Participants completed 200 trials for each of four spatial certainty/reward contingency conditions (.29/fixed, .29/decay, .029/fixed, .029/decay; block order was counterbalanced across participants). Although trial number per condition was substantially lower than in Experiment 1, adjustments to changes in spatial certainty in other cuing studies have shown clear effects with less than half the trial numbers we used here (Pinzmetal et al., 2015; Vossel et al., 2015). We included only these two levels of spatial certainty as we wanted to avoid any possible floor or ceiling effects when testing the influence of reward condition.

We also tested the separate hypothesis that individuals may mentally represent the high and low incentive placeholders differently in terms of their relative value, when their value can be obtained more reliably (i.e. in the fixed reward-condition, relative to the decay reward-condition), and that this may be expressed via physical placement on a linear space. Every 50 trials, participants were instructed to use a mouse to drag the two placeholders wherever they liked on a single line. However, we found no evidence that cue-likelihood influenced placement of the placeholders (p = .96), and this separate aspect of the study is discussed no further. Participants also completed a BIS/BAS questionnaire (Carver & White, 1994) that was used to test a hypothesis for a separate study not reported here.

## Statistical Approach

We followed the same data cleaning procedures as Experiment 1. Again, piecewise linear functions were fit to the data to isolate the trials contaminated by spatial certainty learning effects. The number of trials removed from the start of each block were similar to Experiment 1 (mean = 14.7, sd 8.5).

We also used the same model comparison approach, with the exception that we added the reward condition term to the linear mixed effects models that were fit to the data.

## Results

### RT

Group mean RT data are shown in Figure 4a. Correct responses were approximately 21 ms ± 17 (SE) slower for targets appearing in low versus high incentive value placeholders; valid versus invalid spatial cues speeded RTs on average by 33 ms ±12 (SE); and the decay-reward condition speeded RTs by 54 ms ±17 (SE) on average compared to the fixed-reward condition. Unlike Experiment 1, there was no detectable influence of spatial certainty. (Note: We tested whether this could be attributed to a reduced sensitivity owing to the fewer levels of spatial certainty used in Experiment 2 relative to Experiment 1, by taking the data from Experiment 1 and extracting only the levels of spatial certainty that were used in Experiment 2. A repeated measures ANOVA (incentive value x spatial certainty x cue validity) showed a significant spatial certainty x cue interaction (F(1, 20) = 4.84, MSE = .001, p = .047), suggesting that reducing the levels of spatial certainty is not sufficient to reduce detectability of such an interaction). As in Experiment 1, we identified the most likely model given the data and found that the winning model included main effects of cue validity, incentive value, and reward condition (BF = 2.18E+67 ±.69 %, relative to null model), but did not include an influence of spatial certainty (although see accuracy data). There was good evidence that this was the best model for the data, as it was positively preferred to the next best model, which included an additional incentive value x cue validity interaction term (BF = 4.65 ±2.43 %, see Figure 4b). As spatial certainty was found to interact with cue validity in Experiment 1, we tested the evidence for the winning model against one that also included a spatial certainty x cue validity interaction term. Again, there was positive evidence that the winning model provided a better fit to the data (BF = 8.97 ±1.79%). Collectively, the results show that even when an additive operation is disadvantageous, an additive model is still a better account of the data.

**Figure 4:**
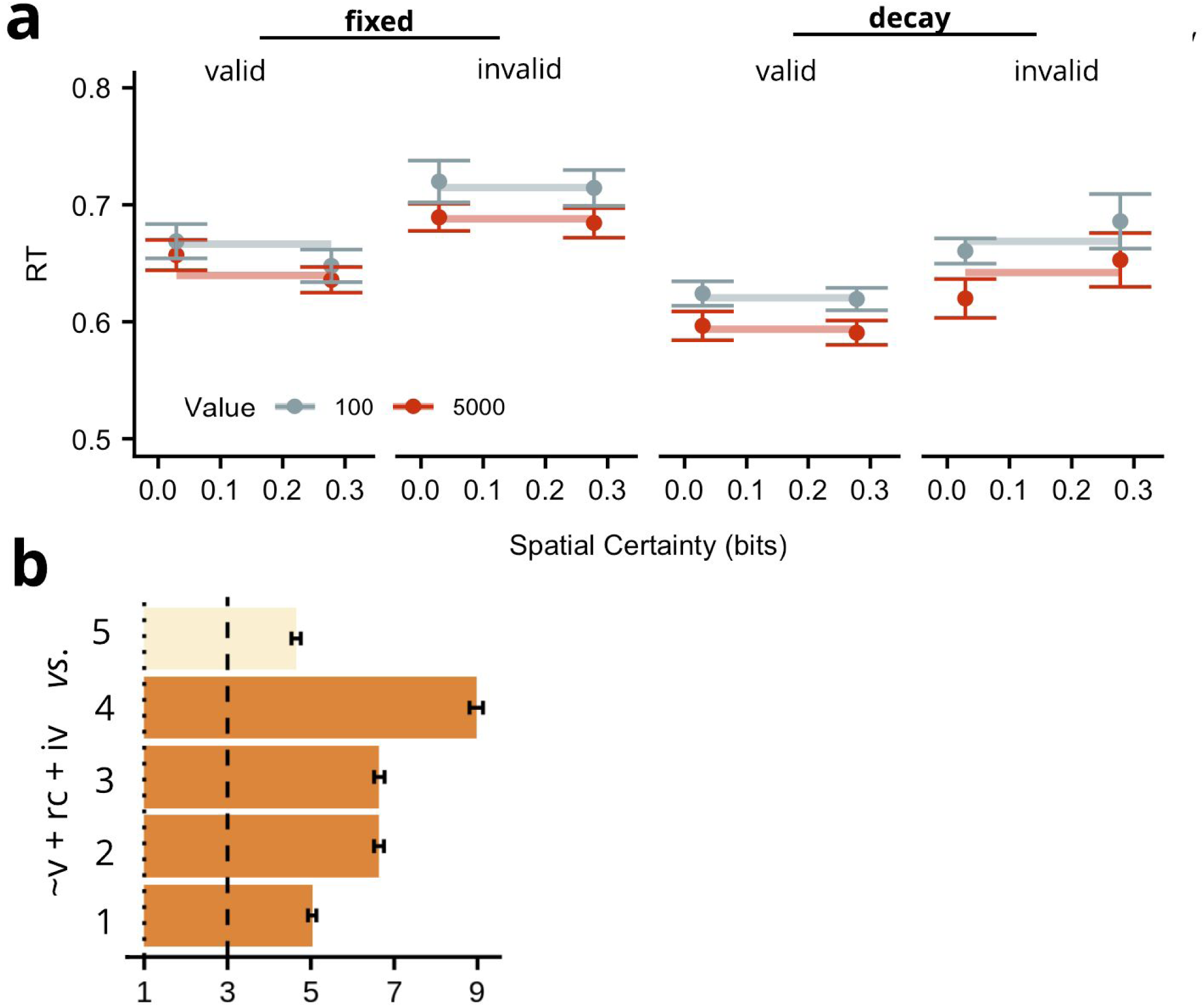
RT Experiment 2. **a**. Observed group mean RTs in ms for targets appearing in high (dark circles) or low (light circles) value placeholders plotted as a function of spatial certainty (x-axis) for valid and invalid spatial cues (panels) for each reward condition. Vertical lines indicate ± 1 within-subject SE. Lines represent fit of the winning model. The winning model involved main effects of incentive value, cue validity, and reward condition. **b**. BFs for the probability of the winning RT model (P[Win]: v + rc + iv), relative to the 5 next best models (Alternative, P[Alt], models, y-axis). The larger the BF value, the stronger evidence for the winning model. Any values lower than 1 (dotted line) support P[Alt]. *The larger the BF value, the stronger the evidence for the winning model*. Any values lower than 1 (dotted line) support P[Alt]; BF values between 3 (dashed line) and 10 constitute moderate evidence for the winning model, values > 10 provide strong evidence. Dark bars indicate P[Alt] only contains an additive influence of incentive value; light bars indicate P[Alt] involves a multiplicative influence of incentive value and either spatial certainty or validity. The Alt RT models were as follows: 1) ∼(rc x iv) + v + iv + rc, 2) ∼AME, 3) ∼(rc x v) + iv + v + rc, 4) ∼(v x sc) + AME, 5) (v x iv) + v + iv + rc. BF = Bayes Factor. valid = cue-validity, sc = spatial certainty, iv = incentive value, rc = reward condition, AME = all main effects.

### Accuracy

Differences in accuracy between valid and invalid trials grew larger as spatial certainty increased (by .15 on average, ±.07 (SE)), suggesting that as participants became slower on invalid trials, they also became less accurate. Furthermore, accuracy performance was slightly higher when targets appeared in high relative to low incentive value placeholders (by approximately .0003, ± .0001 (SE)), showing that participants became very modestly less accurate as they slowed their responses to low value targets. Accuracy was higher for the fixed relative to the decay reward condition (.05, ± .008 (SE)), suggesting that as participants became faster in the delayed reward condition, they also produced a very modest decrease in accuracy. As in Experiment 1, the accuracy data obtained here demonstrate that the validity and incentive value results were not due to a speed-accuracy trade off, whereas the influence of reward condition may indeed be due to such a trade-off (see Figure 5). The best fitting model to account for these data contained a cue-validity x spatial certainty interaction, and main effects of incentive value, spatial certainty and cue validity.

**Figure 5:**
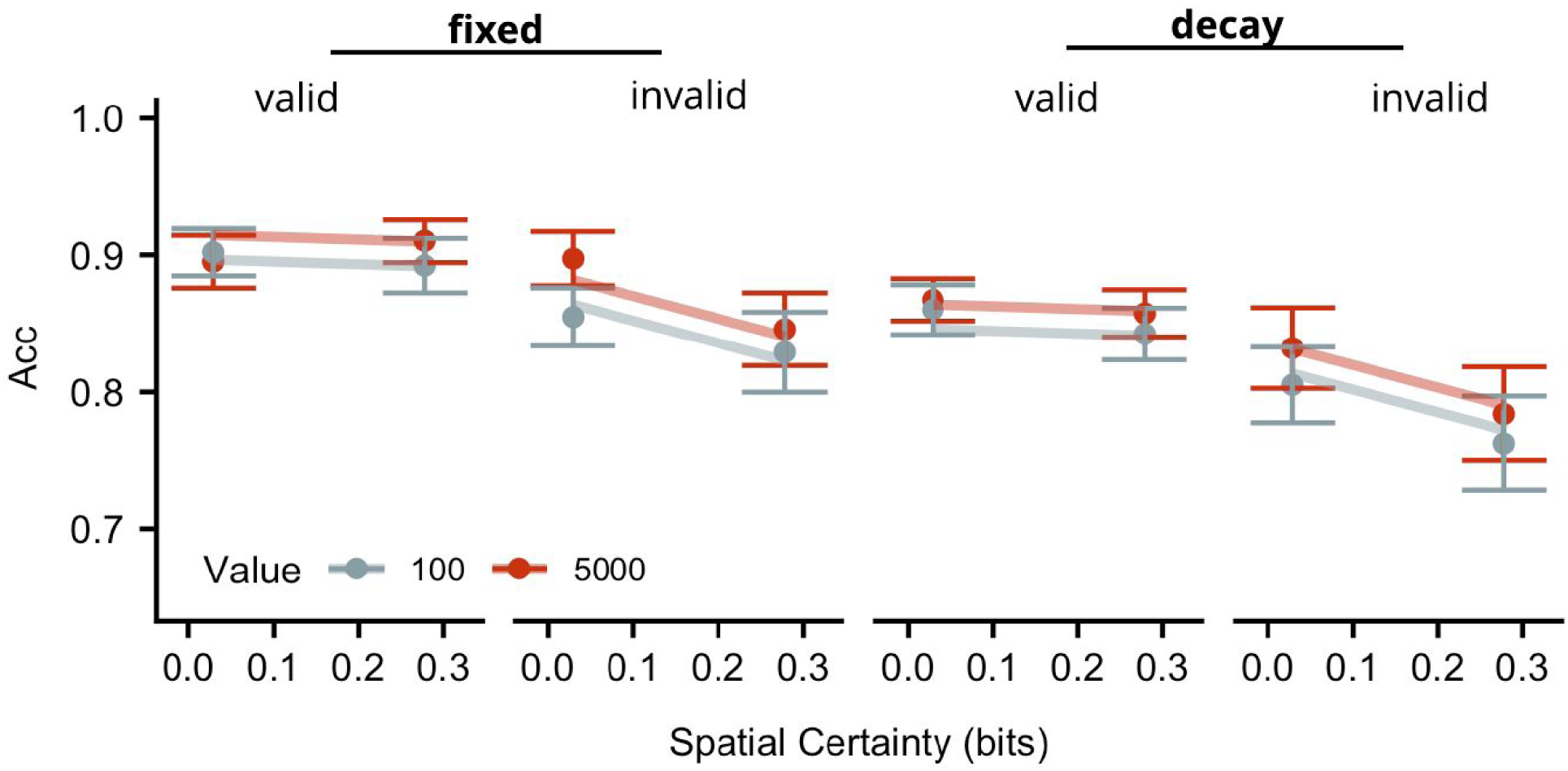
Accuracy data for Experiment 2. Accuracy for targets appearing in high (dark circles) or low (light circles) value placeholders plotted as a function of spatial certainty (x-axis) for valid and invalid spatial cues (panels) for each reward condition. Vertical lines indicate ± 1 within-subject SE. Lines represent fit of the winning model. Acc = accuracy

## General Discussion

Over two experiments we tested whether the additive hypothesis would outperform the expected value account, even under conditions expected to challenge the optimality of additivity. In Experiment 1, we hypothesised that if incentive value and spatial certainty influence a common underlying mechanism, then conditions wherein spatial certainty is trivial to compute (i.e., very high certainty) might best reveal non-additive effects, because these conditions should be minimally taxing to central resources and thus be more likely to enable an influence of incentive value on visual selection. We created this condition by using very high spatial certainty cues and then pitted incentive value and spatial certainty against each other in a spatial-orienting task, where endogenous cues signalled the likely location of upcoming letter targets. Interestingly, an additive influence of incentive value and spatial certainty was observed, even under conditions of very high certainty. Spatial certainty increased the size of the cueing-effect (i.e. the difference in performance between invalidly and validly cued trials), whereas incentive value had a comparable influence on both valid and invalid trial types, across all levels of spatial certainty.

In Experiment 2, we reasoned that if an expected value operation can bias visual selection, then a reward structure that favours a multiplicative weighting of incentive value and spatial certainty may reveal it. We applied an exponential decay function to incentive values at target onset; this ensured that if RTs were driven by multiplicative as opposed to additive weighting, then reward gains accrued by faster RTs under high incentive value/high certainty conditions would outweigh losses incurred by slower RTs under low incentive value/low certainty conditions, relative to RTs predicted by an additive model. Although the influence of spatial certainty manifest differently than in Experiment 1, i.e., by modulating accuracy, rather than RT, we observed that the effect of incentive value remained additive to spatial certainty and to other experimental factors. Again, the findings support the additive hypothesis. However, this interpretation depends on the assumption that participants were able to infer the relationship between response time and rewards. Although participants were both explicitly informed about the decaying reward and were exposed to a sufficient range of the function to theoretically be able to infer the relationship between response time and rewards, an explicit measure of the participant’s understanding of the response-time/reward relationship should be taken in future studies to verify this assumption.

What kind of mechanism or computation could result in a robust additive influence between incentive value and spatial certainty? In concert with recent theoretical and empirical developments suggesting that cognitive control processes are offset by subjective and computational costs of effortful control (Braver, 2012; Yee & Braver, 2018), we believe the current data can be interpreted as reflecting trial by trial adaptations aimed at the conservation of effort. If we assume that the maintenance of the task set, i.e., *a priori* preparedness to identify a data-limited target at two locations, requires energetic resources from the underlying selection mechanism, it is of benefit to the brain to predict conditions where effort can be relaxed, in order to conserve energy expenditure. For example, by learning the energetic range over which target identification mechanisms can be adjusted, to ensure good enough target detection, given the task parameters. According to this view, a cost-benefit analysis could inform how much energy could be saved, given an acceptable decrement to accuracy and response time.

Applied to current context, after the onset of the incentive value cues, selection mechanisms should maintain a steady level of task preparation favouring the high value location for example, by increasing the excitability of neuronal assemblies whose collective receptive fields correspond to detecting lines or edges at that location (Carrasco, 2011; Desimone & Duncan, 1995; Roelfsema et al., 2000; Schmitz & Duncan, 2018), thereby biasing the system towards a stronger response to the upcoming stimulus (Buschman & Kastner, 2015). Concurrently, the excitability of neuronal assemblies directed towards encoding information from the low value location should be relaxed, as the cost of sometimes missing the target at that location, given the energy needed to detect it, should become negligible. Similarly, upon spatial cue onset, preparation of such target detection mechanisms could be further relaxed for the unlikely location, proportionally to how unlikely that location is to possess a target. Importantly, this is performed incrementally to the previous adjustment. This interpretation suggests that the system incrementally updates cost efficient sensory encoding on the basis of incoming information. This interpretation predicts that the degree to which incentive value or spatial certainty can influence performance is dependent upon the range over which preparatory processes can be titrated and still yield acceptable performances. This would account for why, in Experiment 2, under a context with greater pressure on RT performance, we observed an influence of cue validity but not spatial certainty. Presumably, it had become too costly to modulate RT performance by spatial certainty and meet the perceived demands of the task.

The current findings shed further insights into the previously proposed unitary selection mechanism that biases competition between cortical representations of stimuli, in the presence of both incentive and explicit spatial cues (Stanisor et al., 2013). These authors showed that overlapping clusters of macaque V1 neurons were sensitive to both incentive value cues and 100 % valid spatial cues. The authors proposed that the explicit spatial cue served to reweight the relative value between the target and distractor, and that this reweighting was instantiated by a unitary selection mechanism that computes the relative expected value between targets. The current study shows that the spatial certainty derived from explicit cues does not contribute to an expected value operation with the previous incentive values to reweight the relative value between the two items. Rather, an additive influence points to the repeated invocation of a selection mechanism on the basis of updates from separable information sources. As incentive value and spatial certainty appear to have been added, rather than multiplied to the existing output of the selection mechanism, the two sources of information may be transformed into a common representational space, or unit, prior to the enactment of their influence^3^.

An alternate, but not mutually exclusive reason for the independence of incentive value and spatial certainty may be that the two are governed by different learning systems; for example, recent investigations measuring overt eye movements, rather than covert attention shifts, found that incentive values induce associative behaviours (Kim & Anderson, 2019; i.e. participants look at the value cue even when paid not to, Le Pelley et al., 2015), whereas spatial cues may be learned via instrumental conditioning (i.e. participants are more likely to repeat an orienting behaviour that has been reinforced, Kim & Anderson, 2019). Indeed, evidence that visual selection biases remain present when explicit spatial cues (i.e. arrows or words) are non-predictive suggests that these behaviours may be under the control of the habitual learning system (Jiang, 2018; Yin & Knowlton, 2006), which governs learned stimulus-response contingencies. Moreover, in the current experiments, we have taken a snapshot of the relationship between incentive value and spatial certainty at one time point (400 ms SOA), potentially when the system is still in a state of updating. It is possible that a multiplicative relationship between spatial certainty and incentive value emerges later in time, once the information from each cue has undergone further processing. This is a possible avenue of further investigation in future studies.

An additive influence of incentive value on visual selection was also observed by Stankevich and Geng (2014), when value was pitted against the varying probability that a target would appear on one side versus the other, in the absence of explicit spatial cues. A visual comparison of the current data and the data from Stankevich and Geng (2014) also yields some interesting points of difference concerning the influence of spatial certainty in the presence or absence of explicit spatial cues. With the current explicit cues, we observe RTs that are comparable across spatial certainty conditions for valid cues, whereas RTs on invalid trials increase as spatial certainty decreases. We inspected the previous literature and observed that this is a consistent finding across other studies that varied the probabilities of explicit spatial cues in comparable tasks (Lanthier et al., 2015; Vossel et al., 2006), suggesting that this is a replicable phenomenon, rather than the consequence of a ceiling effect or something similar. In contrast, Stankevich and Geng (2014) observed the opposite; RTs decreased as target location became more likely (analogous to valid trials) but remained invariant when directed towards increasingly less likely (invalid) locations. Therefore, our results suggest that the explicit spatial cue we used resulted in preparation towards the cued location that did not vary with the certainty offered by the cue, coupled with a relaxation of preparation towards the invalid location that scaled with certainty. In contrast, non-explicit spatial knowledge appears to cause a strengthening of preparation towards the more likely location, with no concomitant relaxation towards the unlikely location. This suggests that spatial certainty influences visual selection differently dependent on how it is learned. For example, a long history of arrows serving as useful directional cues could motivate a strong response to the directional stimulus, against which other useful behaviours can be adapted. Therefore, it appears that visual selection behaviours adapt to environmental information in relation to the most contextually relevant baseline behaviour.

## Conclusions

Over two experiments, we sought to arbitrate between competing theories for how learned associations pertaining to incentive value and spatial certainty combine to influence visual selection. Specifically, we asked whether this influence was additive (*Additive hypothesis*), or multiplicative (*Expected Value hypothesis*). We tested these hypotheses by pitting incentive values and spatial certainties against one another in a spatial cueing task under conditions expected to challenge the optimality of an additive operation. The data from two experiments support the notion that visual selection mechanisms show independent sensitivity to incentive value and spatial certainty information, and that both information sources are converted to a common representational space, or unit, in order to influence visual selection. We also interpret our results as suggesting that the mechanisms underpinning visual selection dynamically leverage distinct information sources to reflexively conserve effort within a range that allows acceptable performance given the current task parameters.

## Open Practices Statement

The data and materials for both experiments are available online. See: https://github.com/kel-github/attention-value-certainty

## Acknowledgements

K.G. Garner would like to acknowledge Christopher Nolan, Andrew Clouter and Dragan Rangelov for the comments and insightful discussions about this work. This project was supported by grant ES/L000210/1 from ESRC to J Raymond. K.G. Garner is also supported by a Marie Sklodowska Curie Global Fellowship.

The authors declare no competing interests.

## Footnotes

1 https://github.com/kel-github/attention-value-certainty

2 For a demonstration that participants were exposed to a sufficient range of the reward decay function to identify the reward structure, we refer the reader to the online supplement: https://github.com/kel-github/attention-value-certainty/blob/master/code-analysis-and-task/Supplement_IncentiveValueAndSpatialCertainty.pdf

3 An alternate explanation for the observed additive relationship between spatial certainty and incentive value on RT performance is that an expected value operation is being performed by summation on a logarithmic scale. For example, the expected utility of the scene (E(U)) could be computed as follows: 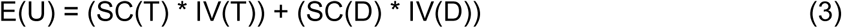 and 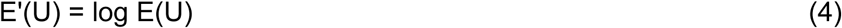 where SC is spatial certainty, IV is incentive value, T is the target location and D is the distractor location. In this case, if the distractor location’s priority was assigned a value of zero, E(U) would reduce to log(SC(T)) + log(IV(T)), i.e., an additive relationship between spatial certainty and incentive value. However, such an account hinges on the notion that the distractor location is somehow assigned a value of zero. This interpretation seems unlikely given that the distractor location has value, and can never be fully ruled out as a potential target location. Moreover, such a computation should produce logarithmic, and not linear relationships between spatial certainty and RTs, when the latter was observed across both Experiments 1 & 2.

